# Mapping the breast tumor microenvironment: proximity analysis reveals spatial relationships between macrophage subtypes and metastasis-initiating cancer cells

**DOI:** 10.1101/2024.03.15.585195

**Authors:** Eloise M. Grasset, Atul Desphande, Jae W. Lee, Yeonju Cho, Sarah M. Shin, Erin M. Coyne, Alexei Hernandez, Xuan Yuan, Zhehao Zhang, Ashley Cimino-Matthews, Andrew J. Ewald, Won Jin Ho

## Abstract

The development of metastasis, responsible for the majority of cancer-related fatalities, is the most dangerous aspect of breast cancer, the predominant malignancy affecting women. We previously identified specific cancer cell populations responsible for metastatic events which are cytokeratin-14 (CK14) and E-cadherin positive in luminal tumors, and E-cadherin and vimentin positive in triple-negative tumors. Since cancer cells evolve within a complex ecosystem comprised of immune cells and stromal cells, we sought to decipher the spatial interactions of these aggressive cancer cell populations within the tumor microenvironment (TME). We used imaging mass cytometry to detect 36 proteins in tumor microarrays containing paired primary and metastatic lesions from luminal or triple-negative breast cancers (TNBC), resulting in a dataset of 1,477,337 annotated cells. Focusing on metastasis-initiating cell populations, we observed close proximity to specific fibroblast and macrophage subtypes, a relationship maintained between primary and metastatic tumors. Notably, high CK14 in luminal cancer cells and high vimentin in TNBC cells correlated with close proximity to specific macrophage subtypes (CD163^int^CD206^int^PDL1^int^HLA-DR^+^ or PDL1^high^ARG1^high^). Our in-depth spatial analysis elucidates that metastasis-initiating cancer cells exhibit with distinct cell populations within the TME, implicating the role of these cell-cell interactions in promoting metastasis.

## Introduction

Breast cancer is the most prevalent malignancy affecting women. Immunohistochemistry analyses have categorized breast cancer into four therapeutic subtypes: luminal A or B tumors, characterized by the expression of hormonal receptors (ER/PR) and varying proliferation rates; human epithelial growth factor receptor 2 (HER2/ERBB2)-amplified tumors; and triple-negative tumors, characterized by the absence of hormonal receptor and HER2 amplification. Although this classification statistically aligns with clinical behavior and treatment responses, it falls short in capturing the complex inter- and intra-tumor diversity unveiled by recent comprehensive genomic and transcriptomic analyses^1–4^.

The most dangerous aspect of cancer is the development of metastasis, responsible for the majority of cancer-related fatalities^5^. To colonize a distant site, cancer cells must first invade their environment to leave the primary tumor and infiltrate the bloodstream or lymphatic system. Once in circulation, cancer cells face an array of challenges that must be surmounted for their survival and subsequent colonization of remote sites. It has been established that only a subset of cancer cells possesses the capability to execute these multiple steps and form metastases^6^.

In the context of luminal breast cancer, our research demonstrated that collective invasion is orchestrated by cancer cells expressing the basal marker cytokeratin 14 (CK14)^7^ and the epithelial marker E-cadherin (ECAD)^8^. Further investigations in preclinical models unveiled that, in the absence of CK14 or ECAD, luminal cancer cells are unable to metastasize^8,9^. Conversely, in triple-negative cancer (TNBC), we established that collective invasion is driven by hybrid E/M cancer cells expressing both the epithelial marker ECAD and the mesenchymal marker vimentin (VIM)^10^. Knocking-down the expression of ECAD or VIM in TNBC cells impeded metastasis formation^8,10^. These studies have shed light on specific cancer cell populations responsible for metastatic events which are the expression of CK14 and ECAD in luminal tumors and the expression of ECAD and VIM in TNBC.

Cancer cells evolve within a complex ecosystem, comprising immune cells and stromal cells, all embedded within a remodeled extracellular matrix. Single-cell analyses have revealed the extent of tumor microenvironment (TME) cellular diversity^3,11^ but have not addressed how these cells are organized in space. However, the interactions of cancer cells within their TME significantly influence cancer cell behavior and response to treatment^12,13^. In a previous study, we established that the aggressive CK14^+^ cells in luminal cancers are more susceptible to natural killer (NK) cells recognition than their CK14^-^ counterparts^14^. This investigation suggests that non-cancerous cells within the TME can interact differentially with cancer cells depending on their states.

The development of multiplexed imaging technologies, including imaging mass cytometry (IMC), enable the classification of cell types and states in the context of spatial relationships. The main limitations of these techniques are the numbers of markers used (usually < 40) and the variation of these markers between studies. However, the resolution of this technology to single cell levels makes it extremely powerful to study cell-cell interactions. Previous studies using IMC confirmed that immune cells are excluded from the tumor mass in most breast cancer tumors and that stromal fibroblasts are often found between cancer cells and immune cells, suggesting that they are agents of immune exclusion^15,16^. They demonstrated that the TME diversity differed markedly among breast cancer subtypes and that genomic alterations in cancer cells influenced the TME composition^15,17^. IMC classification of cancer cells into 18 single-cell pathology subgroups better predict patient outcomes compared to the classical immunohistology classification^16^. However, since the ability to explore specific questions directly relies on what markers have been assayed as a part of the IMC antibody panel, these studies were unable to assess the spatial relationship between cancer cell states and the TME components such as immune cells or stromal cells. Deciphering the spatial interactions of aggressive cancer cell populations with TME components could reveal important relationships supporting cancer cell metastasis initiation and formation.

In this study, our objective was to determine the interactions of distinct cancer cell populations within both primary and metastatic breast cancer TME, with a particular interest in previously characterized aggressive cancer cell populations. To study the cellular composition of breast tumors while preserving spatial context, we used IMC to detect 36 proteins in a tumor microarray (TMA) containing paired primary and metastatic tumors from individual patients with TNBC or luminal breast cancer. Our analysis identified 11 cancer cell states and determined the proportion of these states in both primary and metastatic tumors, with clear differences between luminal and TNBC tumors. We also showed the TME compositions of these tumors, highlighting distinct pattern of immune infiltration between primary and metastatic TNBC tumors. Analysis of cell-cell interaction networks revealed a close proximity between markers of metastasis-initiating cancer cell and specific populations of macrophages and stromal cells.

## Results

### Enumeration of key cancer cell phenotypes and major TME components in primary and matched metastatic breast tumors

To comprehensively quantify the cellular heterogeneity and spatial organization of breast cancer TME, we used the previously established tissue microarray containing cores from hormone-responsive luminal and triple negative breast cancer specimens along with control tissues^18,19^. This cohort is particularly interesting as biospecimens from 26 patients (24 patient samples were available for analysis) were collected from the primary cancer site at the time of initial resection and also from the metastatic sites at the time of recurrence (Supplementary Table 1). The two most common sites of distant metastatic disease available for analysis were brain (10/24) followed by lung (6/24). Other sites of disease included liver, bowel, pancreas, ovary, spine, and pleura. Up to 5 spots per tumors were represented with at least 2 spots per tumor. We analyzed these tumors using imaging mass cytometry (IMC). Briefly, tumors were stained with an antibody panel coupled with rare earth metal reporters. The stained slides were then analyzed by IMC to build high-dimensional images. Finally, images were dissected to determine the phenotype of each cell and its spatial network, which include all cell-cell interactions (Figure 1A).

**Figure 1.**
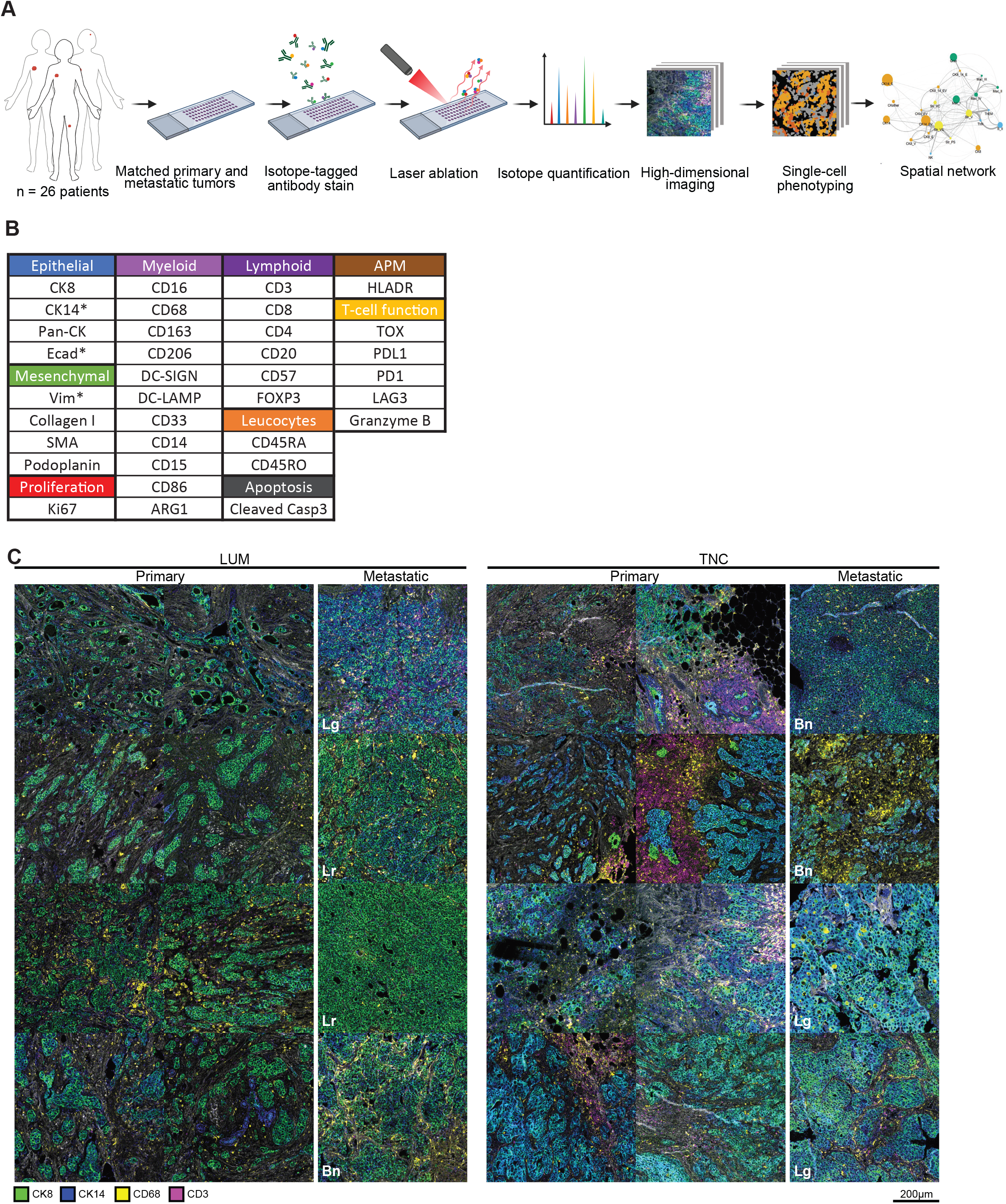
Imaging mass cytometry (IMC) for luminal breast cancers (LUM) and triple-negative breast cancers (TNC). (A) Biospecimens are incorporated into a tissue microarray to be stained together with a cocktail of metal-conjugated antibodies. Tissues are ablated to quantify ion content on a per-pixel (1µm^2^) basis and rendered into images for single-cell segmentation and analysis. Created with BioRender.com (B) Panel of antibodies used. Stars indicate the markers of metastasis-initiated cancer cells (C) Representative IMC results: markers of ductal epithelial cells (CK8 and CK14), macrophages (CD68), and T cells (CD3) are shown for primary and metastatic LUM and TNC. Matching primary and metastatic sites for four unique patients are shown. Abbreviations: Bn, Brain; Lg, Lung; Lr, Liver.

We designed an antibody panel containing 36 markers to distinguish epithelial and mesenchymal cells, myeloid and lymphoid lineages, antigen presentation marker, immune cell function proteins, as well as proliferation (Ki67) and apoptosis (cleaved caspase 3) markers (Figure 1B). Importantly, we included our previously identified markers of metastasis-initiating cancer cell populations: Ecad, K14, and Vim.

Upon acquiring the images, we first verified the quality of the antibody staining, both from the breast cancer specimens (Figure 1C) and control tissues (Figure S1A). These images were then segmented into single cells (Figure S1B). With 337 images, the final dataset represented 353.94mm^2^ tumor area with an average of 14.75mm^2^ per patient. Each image yielded approximately 2,000∼8,000 cells (Figure S1C). The dataset was clustered using FlowSOM^20^ into 50 metaclusters, which were annotated into 11 epithelial, 4 stromal, and 14 key immune cell types based on canonical markers (Figure 2A), totaling 1,477,337 annotated cells. Major immune cell types were identified by predominant expressions of CD3 for T cells (along with CD8^+^ cytotoxic, CD4^+^ helper, and FOXP3^+^ regulatory subtypes), CD20 for B cells, CD57 for NK cells, CD15 for granulocytes, CD68 for macrophages, and DC-SIGN for dendritic cells. Macrophages were further subtyped by varying levels of markers for immunosuppressive states (CD206, CD163) and immunomodulatory functions (programmed death-ligand 1 [PDL1], arginase-I [ARG1], human leukocyte antigen-DR [HLA-DR]). Stromal fibroblast clusters were defined by combined expressions of podoplanin (PDPN), VIM, α-smooth muscle actin (SMA), and collagen I (COL). Epithelial clusters enriched for cytokeratin (CK; CK8, CK14, and pan-CK) expression were distinguished into subtypes by levels of E-cadherin (ECAD) and vimentin (VIM); the relative expression levels of the key markers characterizing these subtypes were consistent in both luminal and TNBC tumors (Figure 2B).

**Figure 2.**
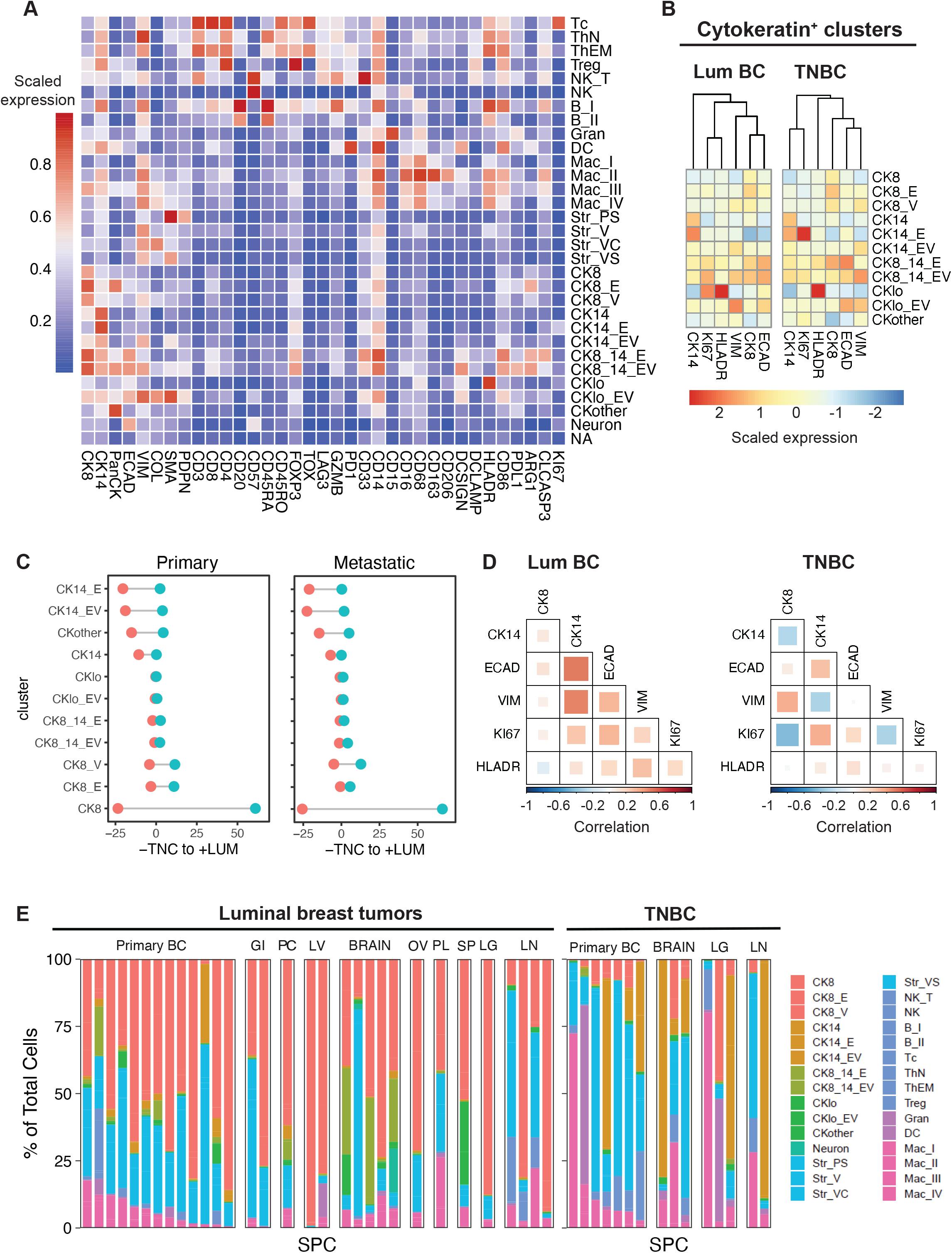
Single-cell profiling with IMC reveals LUM and TNC sub-clusters based on CK8 and CK14 expression and tumor microenvironment (TME) compositions. (A) Results for FlowSOM clustering of IMC data are shown. Scaled expression profile of all markers for each cluster is reflected as heatmap. (B) Scaled expression profiles of selected markers for CK^+^ clusters for LUM and TNC. (C) Differential plot of abundances favoring TNC (negative values) or LUM (positive values) for all CK^+^ clusters. (D) Correlation heatmap of markers relevant to phenotyping CK^+^ clusters. (E) Stacked bar plots of abundances of each cluster faceted by organ site and cancer type from each image. Abbreviations: B, B cells; C, collagen^+^; CK, cytokeratin; DC, dendritic cells; E, e-cadherin; GI, bowel; Gran, granulocytes; LG, lung; LN, lymph node; lo, low expression; LV, liver; LUM, luminal cancer; Mac, macrophage; NA, not assigned; NK, natural killer cells; OV, ovary; PC, pancreas; PL, pleura; PS, PDPN^+^SMA^+^; Str, stromal cells; SP, spine; Tc, cytotoxic T cells; ThEM, effector memory helper T cells; ThN, naïve helper T cells; TNC, triple-negative cancer; Treg, regulatory T cells; V, vimentin

### IMC profiling of cancer cell phenotypes in primary and metastatic lesions from luminal and triple negative breast cancers

We first evaluated the enrichment of CK^+^ subtypes between luminal and TNBC tumors. Consistent with their known profiles^21^, CK8 and CK14 were predominant in luminal cancers and TNBC, respectively, in both primary and metastatic sites (Figure 2C). Next, we investigated the correlations among the markers used to distinguish the CK^+^ clusters and explored their differences between the two breast cancer subtypes (Figure 2D). Notably, we observed a positive correlation between CK14 and VIM expression in luminal tumors but a negative correlation in TNBC tumors. Also, in TNBC tumors specifically, we noticed a negative correlation between CK14 and CK8 as well as between VIM expression and Ki67. Together, these results indicate that the biology of luminal BC and TNBC are distinct and should be analyzed separately.

In luminal primary and metastatic tumors, the most representative cells in terms of cellular composition were epithelial (CK^+^) cells, followed by stromal fibroblasts, macrophages, and the least represented were NK cells, B and T cells (Figure 2E). Among the epithelial cells, the most representative CK^+^ cluster was CK8, followed by CK8_E (CK8^+^ECAD^+^) in primary tumors, or CK8_V (CK8^+^VIM^+^) in metastatic tumors. We also observed an enrichment for CK8_EV (CK8^+^ ECAD^+^VIM^+^) cluster in metastatic tumors. A previous study using a mouse model of luminal cancer demonstrated that chemotherapy induced an EMT in cancer cells that become chemoresistant^22^. Thus, given that all patients with early-stage breast cancer in this cohort received adjuvant chemotherapy (doxorubicin, cyclophosphamide, taxanes, or carboplatinum) prior to disease relapse^19^, it was not unexpected to find enrichment of hybrid E/M clusters, expressing both epithelial (CK8, ECAD) and mesenchymal (VIM) markers, in metastatic tumors compared to matched primary tumors.

Interestingly, 3 out of 5 brain metastatic lesions from luminal cancers exhibited an enrichment for CK8_CK14 clusters, which are not enriched in other metastatic sites comprising bowel (GI), pancreas (PC), liver (LV), ovary (OV), pleura (PL), spine (SP), lungs (LG) and lymph nodes (LN) (Figure 2E). We previously demonstrated that in luminal cancer, CK14 is required for invasion and metastasis seeding. However, we observed a decrease in CK14^+^ cells during lungs metastatic outgrowth in mice^7,9^. Validating these results, we showed that CK14 and CK14_E clusters decreased in luminal metastatic tumors in patients compared to matched primary tumors (Figure S2). This result suggests that the brain microenvironment promotes CK14 expression or maintenance in cancer cells. Further studies analyzing a larger cohort of metastatic patients would be required to verify this hypothesis.

In TNBC, we observed a higher heterogeneity of the TME composition across the primary and metastatic lesions (Figure 2E). The most representative epithelial clusters were CK14_E (CK14^+^ECAD^+^) and CK14_EV (CK14^+^ECAD^+^VIM^+^) in both primary and metastatic tumors, consistent with a basal phenotype (CK14) and the presence of hybrid E/M cells in TNBC primary and metastatic tumors^10^. However, we also detected a significant proportion of CK8 clusters, notably in one lung metastasis, underlining the heterogeneity of these tumors.

Comparing the expression of markers associated with functional states across the CK^+^ clusters in luminal tumors, we noticed that the most proliferative (Ki67-high) clusters were the relatively rare CKlo (low expression of CK) and CK8_14_EV clusters. Conversely, in TNBC the most proliferation was observed in the relatively abundant CK14_E cluster. Additionally, we found in both luminal and TNBC tumors that HLA-DR, an MHC class II cell surface receptor, is enriched in a CKlo cluster that has low levels of ECAD and no VIM (Figure 2B). CKlo cluster is the less abundant cluster in these tumors. MHC class II molecules are important for antigen presentation, e.g., to CD4 T cells, and their presence associated with immune cell infiltration^23^. The general paucity of HLA-DR expression in breast carcinomas has previously been reported^24,25^, implicating one barrier among many that impair effective immune recognition of breast cancers.

### Enumeration of key stromal and immune cell phenotypes in primary and matched metastatic breast TMEs

To decipher the relationships of cancer cell clusters with TME components, we then analyzed the TME composition of these tumors. In primary luminal tumors, as well as in the majority of primary TNBC tumors and their corresponding metastatic lesions, stromal cells (positive for VIM/SMA/PDPN/COL) emerged as the predominant cell type in the TME, constituting a spectrum from 0.67 % to 76.8 % (median 21.1%) of the cells scrutinized per specimen (Figure 3A). Consistent with previous studies that identified multiple carcinoma-associated fibroblast (CAFs) phenotypes in tumors^26–29^, we distinguished four stromal cell populations based on their expression of VIM, COL, SMA, HLADR, and PDPN (Figure 3B). The most representative stromal cell cluster in both luminal and TNBC tumors was Str_VS characterized by the expression of VIM and SMA without PDPN, COL and HLADR. Interestingly, the marker expressions of the four stromal clusters are very similar between luminal and TNBC subtypes, suggesting that these cell states could be transposed between breast cancer subtypes (Figure 3B and C). Comparing stromal cell abundances between primary versus metastatic lesions, we noted a decrease of all stromal clusters in metastasis with significant decreases in Str_PS (PDPN^+^SMA^+^) and Str_VS (Figure S2).

**Figure 3.**
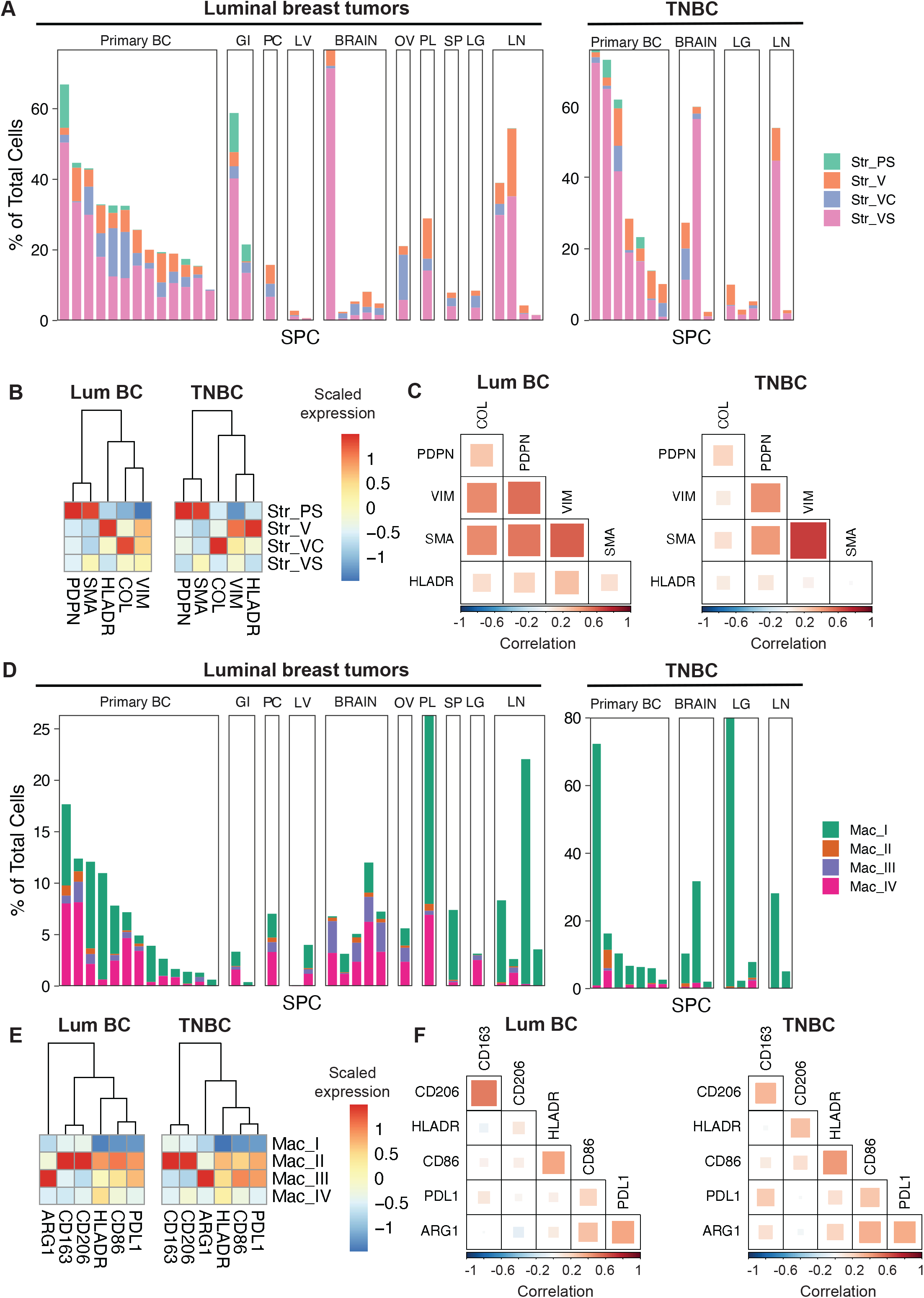
Profiles of stromal and macrophage phenotypes within the TME. (A) Stacked bar plots of cluster abundances of stromal clusters faceted by organ site and cancer type from each image for every case (SPC). Every stacked bar is sorted left-to-right for greatest-to-least abundances of stromal clusters. (B) Scaled expression profiles of select markers for stromal clusters for LUM and TNC. (C) Correlation heatmap of markers relevant to phenotyping stromal clusters. (D) Stacked bar plots of abundances for macrophage clusters faceted by organ site and cancer type from each image for every case (SPC). (E) Scaled expression profiles of select markers for macrophage clusters. (F) Correlation heatmap of markers relevant to phenotyping macrophage clusters. Abbreviations: BC, breast cancer; GI, bowel; LG, lung; LN, lymph node; LUM, luminal cancer; LV, liver; Mac, macrophage; OV, ovary; PC, pancreas; PL, pleura; SP, spine TNC, triple-negative cancer

Among the immune cells, macrophages (CD68^+^ clusters) were by far the most abundant, ranging from 0.05% to 14.1% (median 2.45%) of cells analyzed per specimen (Figure 3D). Four clusters of macrophages were distinguished based on the expression of PDL1, CD86, HLADR, ARG1, CD206, and CD163 (Figure 3E). HLADR and CD86 are usually involved in T cell activation whereas PDL1, ARG1, CD206 and CD163 are markers of M2-polarized macrophages, involved in immunosuppression and anti-inflammatory responses^30^. Strong correlations were observed between PDL1 and ARG1 and also between CD163 and CD206 (Figure 3F). These phenotypic associations were expected given the known involvement of these markers in the immunosuppressive function of macrophages. The most abundant macrophages clusters are Mac_I and Mac_IV in luminal tumors, characterized by low expression of these markers (CD163^int^CD206^int^), and expression of HLADR (CD163^int^CD206^int^PDL1^int^HLA-DR^+^), respectively. In TNBC, Mac_I cluster is the most abundant macrophage subtype in both primary and metastatic lesions, and Mac_IV relatively low (Figure 3D). Interestingly, we observed strong similarities between macrophages clusters in terms of marker expressions in luminal and TNBC tumors (Figures 3E and F). These results indicate the presence of similar macrophage polarizations but with different abundances between the breast cancer subtypes. We did not detect significant differences in macrophages abundances between primary and metastatic lesions (Figure S2).

Overall, other immune cell types including granulocytic cells (CD15^+^; median 0.06%) and B cells (CD20^+^; median 0.27%) were generally rarer in the dataset with T cells (CD3^+^ clusters; median 0.02%) being among the least abundant cell populations especially in luminal cancers (Figure 2E). These results confirmed previous studies characterizing luminal tumors as ‘colder’ than TNBC tumors^31,32^. We did not detect significant difference between primary and metastatic tumors (Figure S2).

### Spatial interactions within the breast TME

To determine how the different cell types are spatially coordinated with one another based on all of the acquired images, we then calculated the average shortest distances between every cell type and visualized all of the cell type clusters as spatial networks (Figure 4). The size of the node indicates the abundance of the cluster and the thickness of the line the proximity, with thicker lines indicating closer clusters.

**Figure 4.**
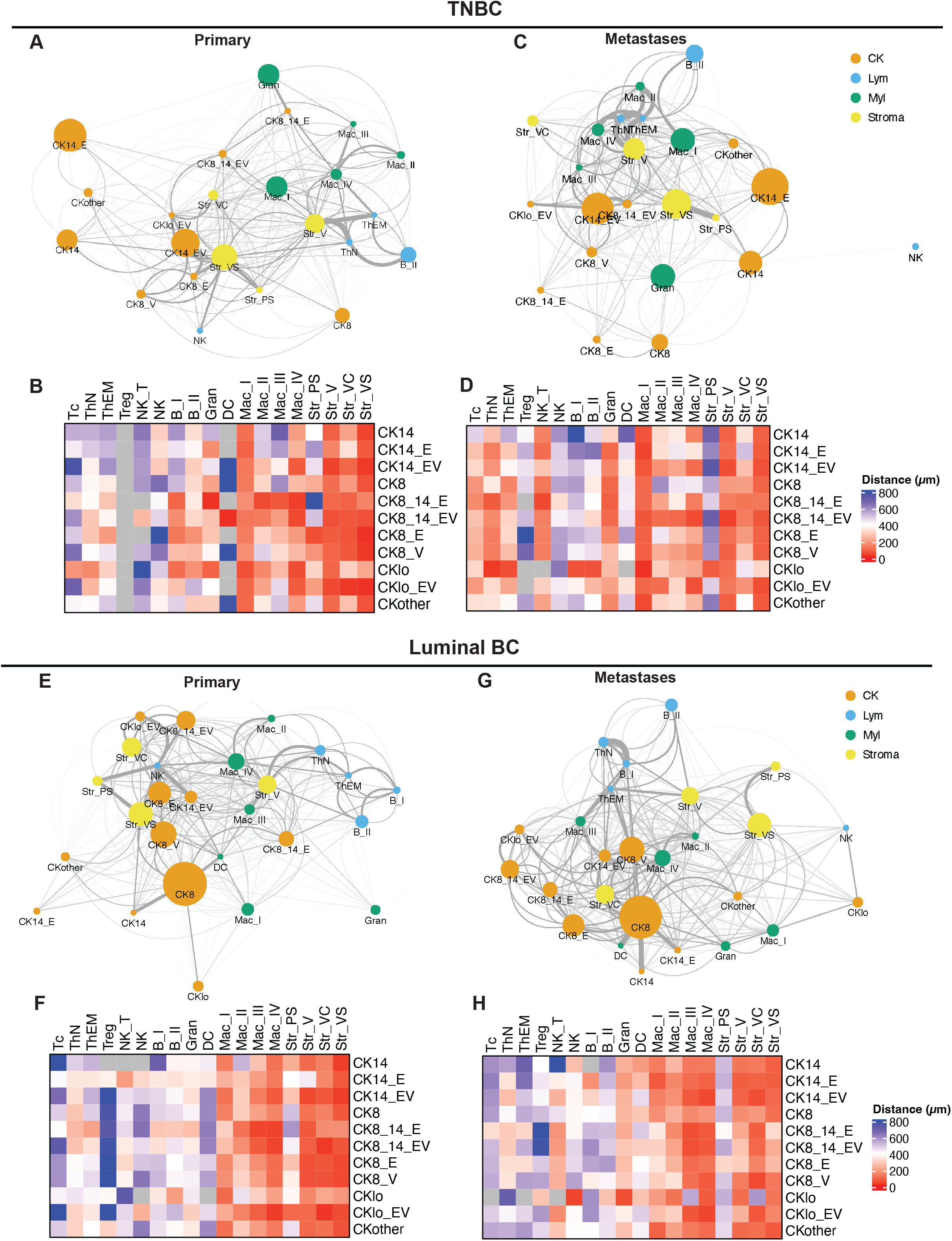
Spatial relationships of CK^+^ and TME cell types. Network visualization plots show distance relationships among clusters based on average shortest distances between cell types for TNBC primary (A), TNBC metastases (C), LUM primary (E) and LUM metastases (G). Node sizes reflect relative abundance of the cell type. Edge thickness positively correlates with shorter distances. Heatmaps of mean distances (µm) between CK+ and TME cell types within each image and then averaged across the entire dataset are shown for TNBC primary (B), TNBC metastases (D), LUM primary (F) and LUM metastases (H). Abbreviations: B, B cells; C, collagen^+^; CK, cytokeratin; DC, dendritic cells; E, e-cadherin; Gran, granulocytes; lo, low expression; Lym, lymphoid; LUM, luminal cancer; Mac, macrophage; Myl, myeloid; NA, not assigned; NK, natural killer cells; PS, PDPN^+^SMA^+^; Str, stromal cells; Tc, cytotoxic T cells; ThEM, effector memory helper T cells; ThN, naïve helper T cells; TNC, triple-negative cancer; Treg, regulatory T cells; V, vimentin

In TNBC primary tumors, CK^+^ cells are closer to each other and stromal cluster VS, with the exception of CK8 cells being isolated (Figure 4A). We noted that T cells and B cells are excluded from cancer cells by Str_V stromal cells and Mac_IV macrophages, both expressing HLADR. It has previously been shown that a subpopulation of CAF named CAF S1 can attract and retain CD4^+^CD25^+^ T cells promoting their differentiation to Tregs, and therefore creating an immunosuppressive environment^28^. While we did not use markers specific to CAF S1 and only captured a rare presence of Tregs, i.e., insufficient for analysis, the spatial distribution of Str_V and the exclusion of T cells suggest the possibility that these stromal cells blocked T cells from infiltrating the tumor. We also observed that hybrid E/M cancer cells are closer to stromal cells and macrophages than other cancer cell clusters. To better visualize cancer cell clusters proximity with the different component of the TME, we represented the distances using heatmaps, with close proximity being represented in red, and larger distance in blue (Figure 4B). Focusing on the hybrid E/M cancer cell clusters, which we know contribute critically to metastasis formation in TNBC^10^, we observed their close proximity to the different stromal clusters, with the exception of Str_PS. The macrophage cluster labeled as Mac_I is situated in close proximity to all cancer cell clusters, while Mac_IV is closer to hybrid E/M clusters compared to other CK^+^ clusters.

Interestingly, we observed a different spatial distribution of the TME components in TNBC metastases (Figure 4C), with a closer proximity of cancer cell clusters to T cells, macrophages, NK cells, and granulocytes (Gran, CD15^+^) (Figure 4D). Despite this close proximity, T cells and B cells remained closer to macrophage clusters and stromal cluster Str_V than cancer cells. Infiltration of specific immune cells can promote metastasis formation and outgrowth. For instance, we previously showed that cancer cells educate NK cells to promote metastatic seeding and outgrowth^14^. Furthermore, in a mouse model of TNBC, neutrophils were shown to be recruited at the metastatic site to promote cancer cell metastasis formation and chemoresistance^33,34^. The enrichment of granulocytes in TNBC metastasis is compatible with the presence of neutrophils, even though specific neutrophils markers should be used to validate this result. Despite these pro-metastatic roles described in literature, the close proximity of immune cells with cancer cells in TNBC metastasis suggest that immunotherapies would be more effective in the metastatic disease. Indeed, one of the main requirements for effective immunotherapies is the contact of immune cells with cancer cells to allow immune cells killing of cancer cells.

In luminal primary tumors, we observed a similar spatial exclusion of T cells and B cells by Str_V stromal cells and Mac_IV macrophages (Figure 4E), suggesting that these clusters have a comparable function in TNBC and luminal tumors. We also observed that cancer cells are in close proximity with stromal clusters and macrophages (Figure 4F). Contrary to TNBC primary tumors, we noted the presence of NK cells in close proximity with cancer cells, particularly the CK14_E cluster, characterizing the metastasis-initiating cancer cells. However, the average distance between NK cells and cancer cells remains above 200 µm, implying that direct contact is relatively limited (Figure 4F).

We observed differences in TME organization between metastatic lesions and primary tumors (Figures 4E and 4G) even though the distance between cancer cells and TME components was similar (Figures 4F and 4H). Notably, we observed an exclusion of the stromal clusters Str_VS, Str_V and Str_PS to the periphery of the metastases. The only stromal cluster infiltrated in the metastatic tumor was Str_VC, characterized by high expression of COL. The NK clusters were also excluded to the metastatic lesion periphery. However, macrophages remained in close proximity with cancer cells, notably with Mac_III (ARG1^hi^CD163^int^CD206^int^HLADR^+^) and Mac_IV intermixing with the different cancer cell clusters.

These results revealed conserved spatial interactions of the TME in TNBC and luminal tumors, including a spatial exclusion of T cells and B cells by the stromal cluster Str_V and macrophage Mac_IV in primary tumors, or by Str_V in metastatic lesions. Interestingly, the metastasis-initiating cancer cells, characterized by the expression of ECAD and VIM in TNBC or ECAD and CK14 in luminal tumors, appeared closer to macrophage clusters compared to other cancer cell clusters.

### Metastasis-initiated cancer cells are localized in proximity with macrophages

To verify the proximity of metastasis-initiated cancer cells with macrophages, we then plotted the expression of ECAD and VIM in cancer cells with the proximity with the distinct macrophage clusters in TNBC (Figure 5A). This analysis revealed a correlation between VIM expression and proximity with Mac_III and Mac_IV. In visual verification of this spatial relationship, we confirmed the close proximity of macrophage subtypes and VIM-high CK^+^ cells in our acquired images for TNBC (S3A, S3B). These two macrophage subtypes represent a small fraction of the tumor cells with abundance varying between less than 1∼5 %. Therefore, their proximity with aggressive cancer cell clusters suggests a role of these macrophages in shaping cancer behavior.

**Figure 5.**
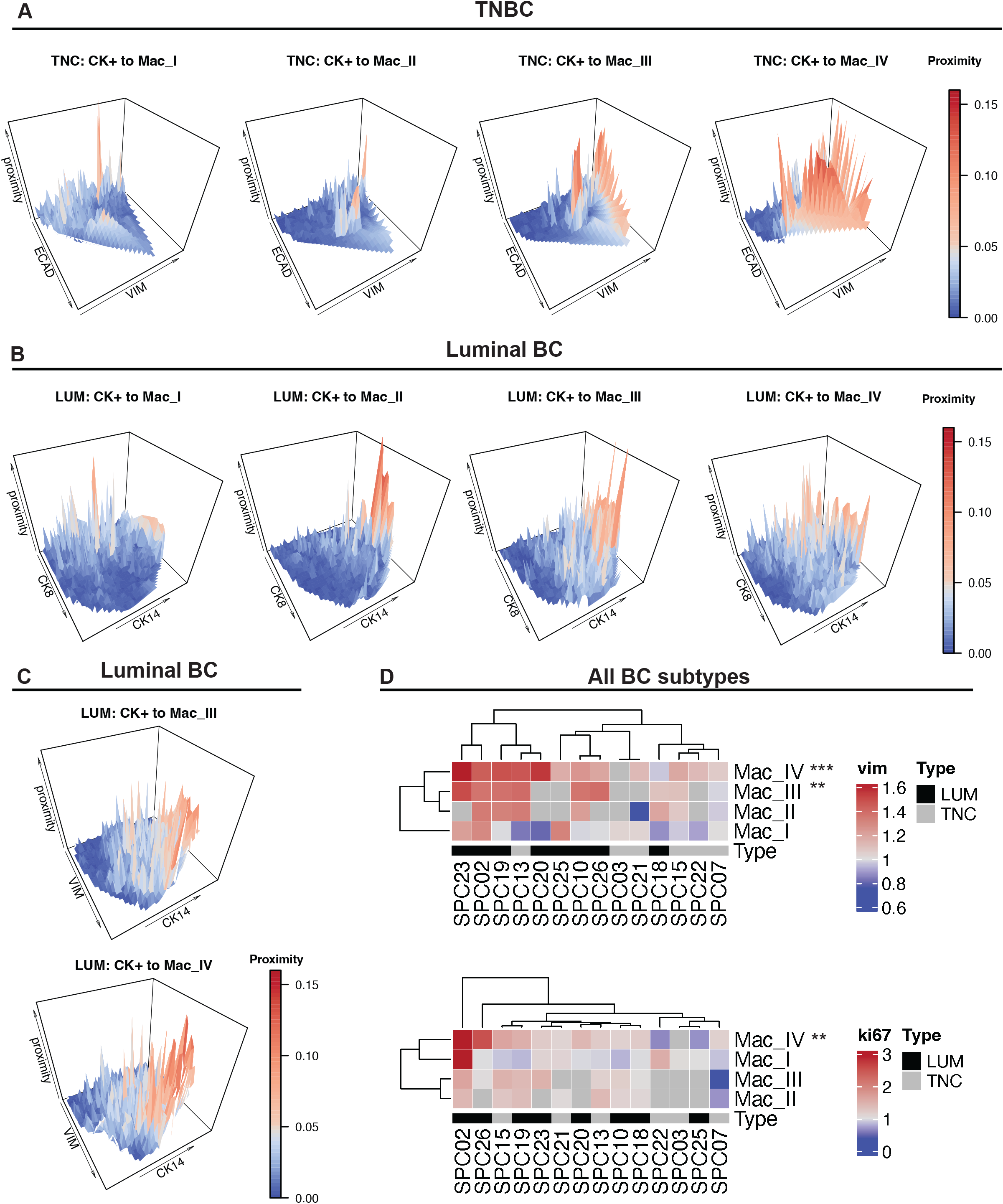
CK+ cells in close proximity with specific macrophage clusters express higher levels of markers associated with metastasis-initiating cancer cells. (A) 3D visualizations of CK^+^ cells in TNBC rendered into peaks for proximity to macrophages (z-axis and scaled color gradient) against their expression of VIM (x-axis) and ECAD (y-axis). (B) Similar 3D visualizations of CK^+^ cells in LUM for proximity to macrophages (z-axis) against their expression of CK14 (x-axis) and CK8 (y-axis). (C) Similar 3D visualizations of CK^+^ cells in LUM against CK14 (x-axis) and VIM (y-axis). (D) Heatmaps for fold-differences in the expression of (top) vimentin and (bottom) Ki-67 by CK^+^ cells that are “close” (<50µm) over “far” (≥50µm) with respect to each macrophage cluster. Expression of vimentin and Ki-67 were compared between close and far CK+ cells using Wilcoxon matched-pairs rank sum test. **p<0.01, ***p<0.005. Abbreviations: BC, breast cancer; ECAD, e-cadherin; LUM, luminal; SPC, specimen/case number; TNC, triple-negative; VIM, vimentin

Interestingly, we observed a similar correlation between cancer cell proximity with macrophages clusters Mac_II, Mac_III, and Mac_IV and CK14 expression in luminal cancer cells (Figure 5B), which characterized the metastasis-initiating cancer cell population. As we noticed a strong correlation between CK14 expression and VIM expression in luminal tumors (Figure 2B), we verified the proximity between macrophage clusters and VIM expression in luminal cancer cells as well (Figure S3A, S3B). Similar to TNBC tumors, in luminal breast cancer lesions, we observed a strong correlation between VIM expression and proximity with macrophages cluster Mac_III and Mac_IV. As expected, when looking at CK14 vs. VIM, the double positive quadrant had higher proximity to macrophages (Figure 5C). We confirmed the close proximity of macrophage subtypes and VIM-high CK^+^ cells in our acquired images for luminal tumors (Figure S3B).

To further validate the close proximity of cancer cells expressing VIM with macrophages, we then compared the expression of VIM in CK^+^ cells that were close (< 50 µm) to macrophage subtypes versus CK^+^ cells that were far. We showed that the expression of VIM was significantly higher for CK^+^ cells that were close to Mac_III and Mac_IV subtypes (Figure 5D). This suggested that distinct subpopulations of macrophages may be driving cancer behavior.

Another behavior of cancer cells that is associated with aggressivity is proliferation. Indeed, high Ki67 index in patient is associated with poor outcome and metastatic recurrence^35^. We measured the expression of Ki67 in cancer cells that were close (< 50 µm) to macrophage subtypes versus CK^+^ cells that were far. Our analysis demonstrated that the proximity with Mac_IV is associated with a higher Ki67 expression (Figure 5D).

While the markers in the IMC panel used could not further distinguish Mac_IV from the other macrophage subtypes other than its intermediate expression of markers involved in immune regulation, e.g., PD-L1 and ARG1, these distance relationships with the expression of VIM and Ki-67 on cancer cells implicated the importance of the Mac_IV cluster on breast cancer cell behavior.

Even though recent seminal studies have spatially profiled breast cancers using IMC, ours is the first study to date that has incorporated the simultaneous analysis of multiple macrophage subtypes based on a combination of CD68, PD-L1, ARG1, HLA-DR, CD86, CD163, and CD206 along with markers of metastasis-initiating cancer cells including ECAD, VIM and CK14. Furthermore, while small in size, this unique cohort of biospecimens have been procured at the time of primary resection and metastatic recurrence and thus are matched by primary and metastatic sites for each patient. This enabled our rigorous analysis of spatial coordination among CK^+^ cells and macrophage subtypes that correlate with differential expression of VIM and Ki-67.

## FIGURE LEGENDS

**Supplementary Figure 1.**
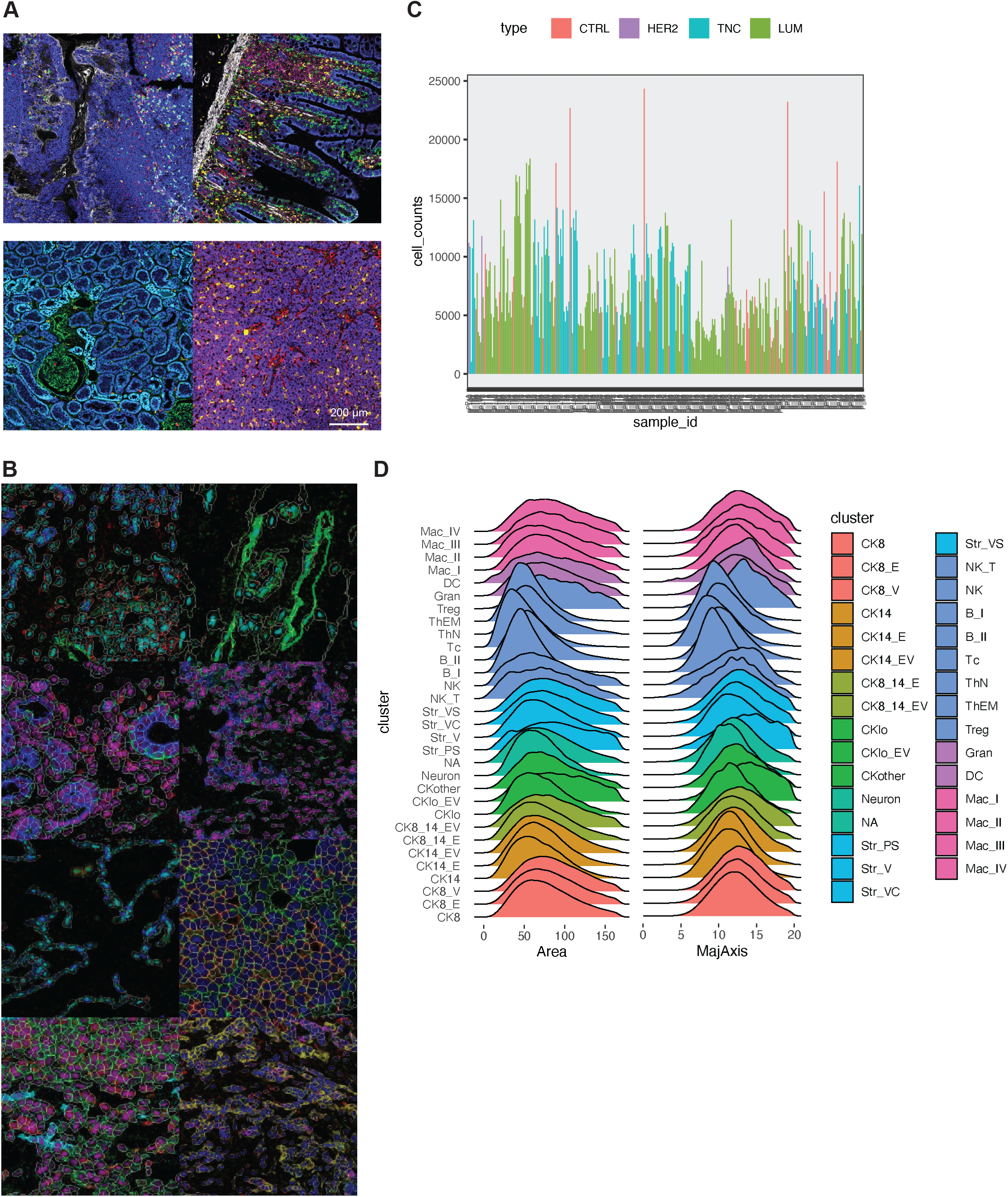
IMC workflow results in robust single-cell data. (A) Representative imaging results for markers for detailed phenotyping validated by control tissues embedded in the same tissue microarray (clockwise from upper left panel: tonsil, intestinal mucosa, liver, kidney). Scale bar is 200 µm. (B) Verification of high quality single-cell segmentation by superimposing segmentation masks over a diverse set of cellular morphologies. 8 representative images are shown at varying scales for the purpose of visualizing the segmentation quality in detail. (C) Total cell counts per region of interest (ROI) yielded by image segmentation. (D) Distribution of key geometric parameters for the segmented cell types, total cell area (µm^2^) and major axis length (MajAxis, µm), are shown. Abbreviations: B, B cells; C, collagen^+^; CK, cytokeratin; CTRL, control tissues; DC, dendritic cells; E, e-cadherin; Gran, granulocytes; HER2, HER2^+^ cancer; lo, low expression; LUM, luminal cancer; Mac, macrophage; NA, not assigned; NK, natural killer cells; PS, PDPN^+^SMA^+^; Str, stromal cells; Tc, cytotoxic T cells; ThEM, effector memory helper T cells; ThN, naïve helper T cells; TNC, triple-negative cancer; Treg, regulatory T cells; V, vimentin

**Supplementary Figure 2.**
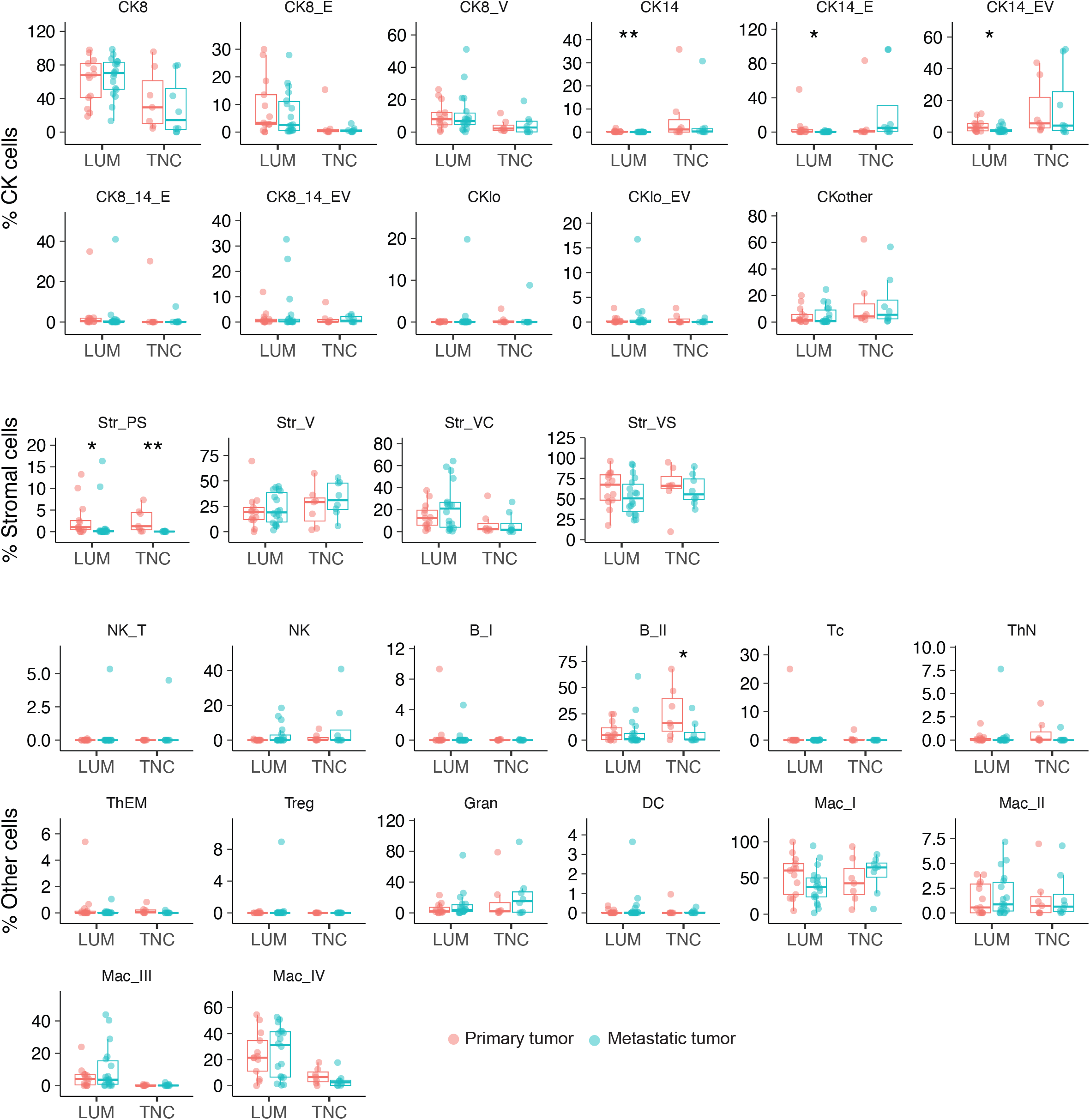
Differences in cluster abundances. Abundances (% cells) of every cell type were compared between primary (red) and metastatic site (turquoise) from LUM and TNC for matched primary and metastatic sites. Differences were compared using Wilcoxon test. *<0.05, **p<0.01. Abbreviations: B, B cells; C, collagen^+^; CK, cytokeratin; DC, dendritic cells; E, e-cadherin; Gran, granulocytes; lo, low expression; LUM, luminal cancer; Mac, macrophage; NK, natural killer cells; PS, PDPN^+^SMA^+^; Str, stromal cells; Tc, cytotoxic T cells; ThEM, effector memory helper T cells; ThN, naïve helper T cells; TNC, triple-negative cancer; Treg, regulatory T cells; V, vimentin

**Supplementary Figure 3.**
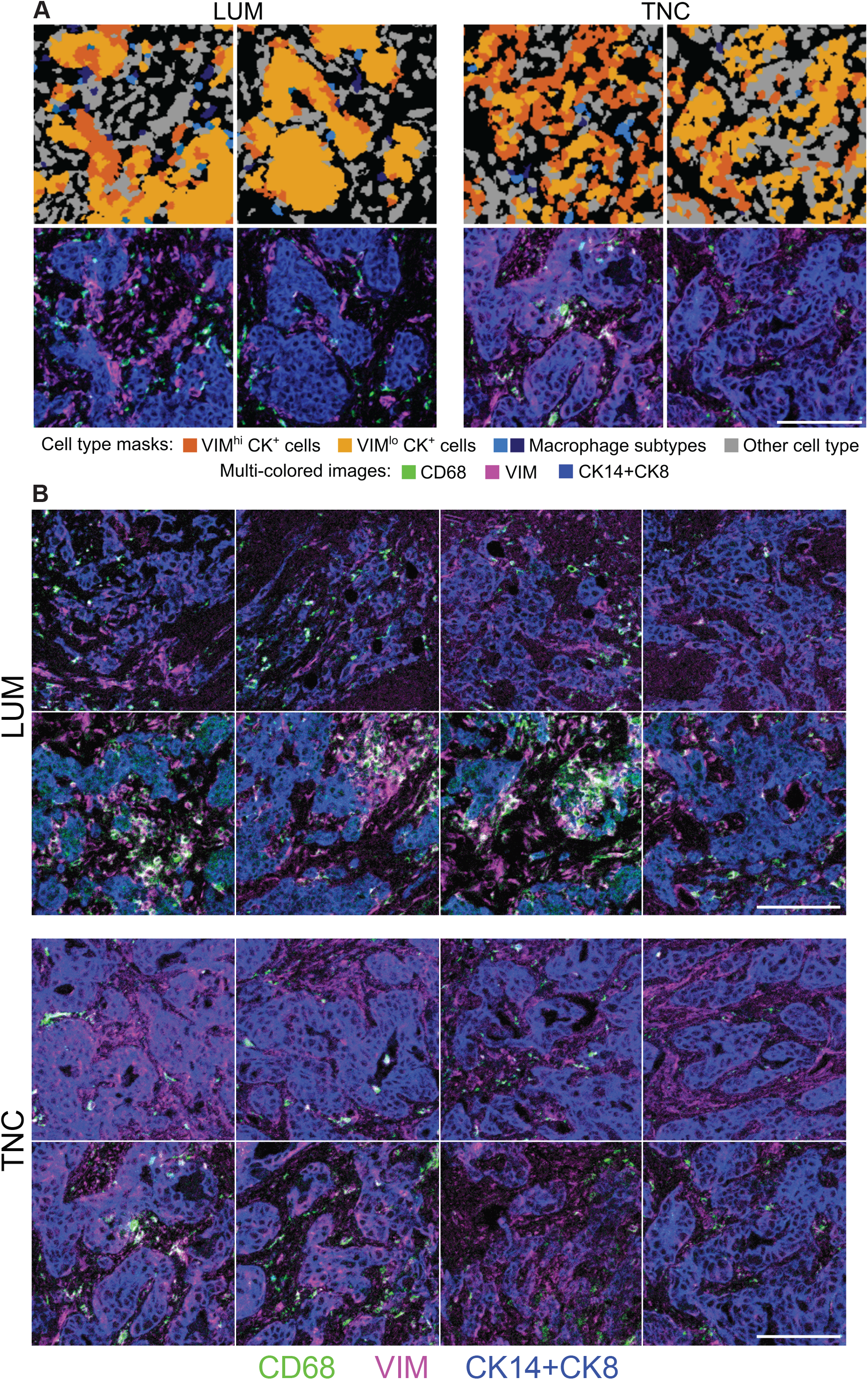
Tumor (CK+) cells in close proximity with macrophage clusters express higher levels of vimentin. (A) Top panels show cell type mapping by colored masks based on cluster annotations, and bottom panels show imaging results from corresponding regions. Scale bar: 100µm. (B) Additional representative images for proximity between macrophages and vimentin-high CK^+^ tumor cells. Scale bar: 100µm. Abbreviations: CK, cytokeratin; lo, low expression; LUM, luminal cancer; TNC, triple-negative cancer; V, vimentin.

## Methods

### Tissue microarrays

Tissue microarrays were generated by a pathologist (ACM) selecting appropriate regions of interest (ROIs) from archived formalin-fixed paraffin embedded tissue blocks from both primary and metastatic breast cancer specimens, followed by extraction of tissue cores with a biopsy needle and arranging them in a recipient block using a tissue arrayer. The block was then baked, and sections were cut for analysis. Studies were conducted in accordance with the 1996 Declaration of Helsinki and were approved by the institutional review board (IRB) at the Johns Hopkins Hospital.

### IMC staining

IMC was conducted on the tissue microarray slides. Slides were baked for 2 hours at 60°C. Dewaxing in xylene, rehydration in alcohol gradient, and heat-mediated antigen retrieval in Antigen Retrieval Agent pH9 (Agilent S2367) at a sub-boiling temperature for 30 minutes were carried out. The slides were blocked with 3% BSA in Maxpar (metal-free) PBS for 45 minutes at room temperature and subsequently labeled with a panel of metal-tagged antibodies targeting a set of antigens including cytokeratins and immune cell markers (Supplementary Table 2) at 4°C overnight. Custom antibodies were conjugated in-house, diluted to a concentration of 0.25mg/mL to 0.5mg/mL, then titrated empirically. For nuclear counterstaining, Cell-ID Intercalator-Ir (Standard Biotools) diluted at 1:400 in Maxpar PBS was applied for 30 minutes at room temperature. For tissue counterstaining, ruthenium tetroxide 0.5% (Electron Microscopy Sciences PN 20700-05) diluted at 1:2000 in Maxpar PBS was applied for 3 minutes at room temperature followed by agitated washing with Maxpar water. After drying the slide, the slide was loaded in the Hyperion Imaging System (Standard Biotools) for image acquisition at the Johns Hopkins Mass Cytometry Facility. Upon selecting all of the cores as ROIs, laser ablation was performed to ionize the metal tags and the resulting ions were detected and rendered into images.

### IMC preprocessing and analysis

Every image was evaluated for data quality and technical artifacts such as ripped tissue and lack of tumor represented. Representative multi-colored images were optimized and exported using MCD Viewer 1.0.560.2 (Standard Biotools). 5 of 342 ROIs of poor quality were excluded from analysis. Image segmentation was carried out using a set of software tools within a standardized pipeline^36^ based on the following softwares: Jupyter Notebook, CellProfiler 3.1.9^37^ Ilastik 1.3.3post3^38^, and HistoCAT 1.76^39^. Stacks of images suitable for all downstream steps in the pipeline were first created using CellProfiler. Pixel-based classification was trained via Ilastik using a combination of nuclear, cytoplasmic, and membranous markers. Upon generating trained probability maps, segmentation masks for all of the images were created using CellProfiler. Quality of the segmentation for all of the images were verified visually and single cell datasets were exported using HistoCAT. All data was then loaded into R 4.2.2 for clustering, abundance, and spatial analyses. Clustering was carried out using *FlowSOM*^20^ and the resulting metaclusters were annotated into cell types based on canonical markers. Spatial calculations were done with *spatstat*, and visualizations were done using *ggplot* and *igraph* packages as well as *plot3D* and *akima* for 3D renders in R.

### Statistics

Statistical tests were performed using R 4.2.2 and GraphPad Prism 9. For exploring abundances of cell types between primary and metastatic sites, Wilcoxon test was used. Comparing VIM and Ki67 expression (per-cell) in CK^+^ cell clusters when close (3∼4 cell range, i.e., <50um) vs. far from the different macrophage clusters was performed across all patients using Wilcoxon matched pairs rank sum.

### Code and data availability

Analysis scripts are available at the following repository: github.com/wjhlab/BreastMetIMC. IMC data are available at the following DOI: 10.5281/zenodo.10688895.

## References

1. Thennavan, A. et al. Molecular analysis of TCGA breast cancer histologic types. Cell Genomics 1, 100067 (2021).

2. Wu, S. Z. et al. A single-cell and spatially resolved atlas of human breast cancers. Nat Genet 53, 1334–1347 (2021).

3. Chung, W. et al. Single-cell RNA-seq enables comprehensive tumour and immune cell profiling in primary breast cancer. Nat Commun 8, 15081 (2017).

4. Lehmann, B. D. et al. Multi-omics analysis identifies therapeutic vulnerabilities in triple-negative breast cancer subtypes. Nat Commun 12, 6276 (2021).

5. Siegel, R. L., Miller, K. D., Fuchs, H. E. & Jemal, A. Cancer statistics, 2022. CA: A Cancer Journal for Clinicians 72, 7–33 (2022).

6. Massagué, J. & Ganesh, K. Metastasis-Initiating Cells and Ecosystems. Cancer Discov 11, 971–994 (2021).

7. Cheung, K. J., Gabrielson, E., Werb, Z. & Ewald, A. J. Collective Invasion in Breast Cancer Requires a Conserved Basal Epithelial Program. Cell 155, 1639–1651 (2013).

8. Padmanaban, V. et al. E-cadherin is required for metastasis in multiple models of breast cancer. Nature 1–6 (2019) doi:10.1038/s41586-019-1526-3.

9. Cheung, K. J., et al. Polyclonal breast cancer metastases arise from collective dissemination of keratin 14-expressing tumor cell clusters. PNAS 113, E854–E863 (2016).

10. Grasset, E. M. et al. Triple-negative breast cancer metastasis involves complex epithelial-mesenchymal transition dynamics and requires vimentin. Science Translational Medicine 14, eabn7571 (2022).

11. Azizi, E. et al. Single-Cell Map of Diverse Immune Phenotypes in the Breast Tumor Microenvironment. Cell 174, 1293–1308.e36 (2018).

12. Wu, P. et al. Adaptive Mechanisms of Tumor Therapy Resistance Driven by Tumor Microenvironment. Frontiers in Cell and Developmental Biology 9, (2021).

13. de Visser, K. E. & Joyce, J. A. The evolving tumor microenvironment: From cancer initiation to metastatic outgrowth. Cancer Cell 41, 374–403 (2023).

14. Chan, I. S. et al. Cancer cells educate natural killer cells to a metastasis-promoting cell state. J Cell Biol 219, e202001134 (2020).

15. Danenberg, E. et al. Breast tumor microenvironment structures are associated with genomic features and clinical outcome. Nat Genet 54, 660–669 (2022).

16. Jackson, H. W. et al. The single-cell pathology landscape of breast cancer. Nature 578, 615–620 (2020).

17. Ali, H. R. et al. Imaging mass cytometry and multiplatform genomics define the phenogenomic landscape of breast cancer. Nat Cancer 1, 163–175 (2020).

18. Cimino-Mathews, A. et al. Androgen Receptor Expression Is Usually Maintained in Initial Surgically-Resected Breast Cancer Metastases, But Often Lost in End-stage Metastases Found at Autopsy. Hum Pathol 43, 1003–1011 (2012).

19. Cimino-Mathews, A., Ye, X., Meeker, A., Argani, P. & Emens, L. A. Metastatic triple-negative breast cancers at first relapse have fewer tumor-infiltrating lymphocytes than their matched primary breast tumors: a pilot study. Hum Pathol 44, 2055–2063 (2013).

20. Van Gassen, S. et al. FlowSOM: Using self-organizing maps for visualization and interpretation of cytometry data. Cytometry A 87, 636–645 (2015).

21. Gusterson, B. A., Ross, D. T., Heath, V. J. & Stein, T. Basal cytokeratins and their relationship to the cellular origin and functional classification of breast cancer. Breast Cancer Res 7, 143–148 (2005).

22. Fischer, K. R. et al. Epithelial-to-mesenchymal transition is not required for lung metastasis but contributes to chemoresistance. Nature 527, 472–476 (2015).

23. Oldford, S. A., Robb, J. D., Watson, P. H. & Drover, S. HLA-DRB alleles are differentially expressed by tumor cells in breast carcinoma. Int J Cancer 112, 399–406 (2004).

24. Wintzer, H. O., Benzing, M. & von Kleist, S. Lacking prognostic significance of beta 2-microglobulin, MHC class I and class II antigen expression in breast carcinomas. Br J Cancer 62, 289–295 (1990).

25. Concha, A. et al. Different patterns of HLA-DR antigen expression in normal epithelium, hyperplastic and neoplastic malignant lesions of the breast. Eur J Immunogenet 22, 299– 310 (1995).

26. Friedman, G. et al. Cancer-associated fibroblast compositions change with breast cancer progression linking the ratio of S100A4+ and PDPN+ CAFs to clinical outcome. Nat Cancer 1, 692–708 (2020).

27. Öhlund, D. et al. Distinct populations of inflammatory fibroblasts and myofibroblasts in pancreatic cancer. J Exp Med 214, 579–596 (2017).

28. Costa, A. et al. Fibroblast Heterogeneity and Immunosuppressive Environment in Human Breast Cancer. Cancer Cell 33, 463–479.e10 (2018).

29. Wu, S. Z. et al. Stromal cell diversity associated with immune evasion in human triple-negative breast cancer. EMBO J 39, e104063 (2020).

30. Mantovani, A., Allavena, P., Marchesi, F. & Garlanda, C. Macrophages as tools and targets in cancer therapy. Nat Rev Drug Discov 21, 799–820 (2022).

31. Denkert, C. et al. Tumour-infiltrating lymphocytes and prognosis in different subtypes of breast cancer: a pooled analysis of 3771 patients treated with neoadjuvant therapy. Lancet Oncol 19, 40–50 (2018).

32. Stanton, S. E., Adams, S. & Disis, M. L. Variation in the Incidence and Magnitude of Tumor-Infiltrating Lymphocytes in Breast Cancer Subtypes: A Systematic Review. JAMA Oncol 2, 1354–1360 (2016).

33. Park, J. et al. Cancer cells induce metastasis-supporting neutrophil extracellular DNA traps. Sci Transl Med 8, 361ra138 (2016).

34. Mousset, A. et al. Neutrophil extracellular traps formed during chemotherapy confer treatment resistance via TGF-β activation. Cancer Cell 41, 757–775.e10 (2023).

35. Soliman, N. A. & Yussif, S. M. Ki-67 as a prognostic marker according to breast cancer molecular subtype. Cancer Biol Med 13, 496–504 (2016).

36. Windhager, J. et al. An end-to-end workflow for multiplexed image processing and analysis. Nat Protoc 18, 3565–3613 (2023).

37. Carpenter, A. E. et al. CellProfiler: image analysis software for identifying and quantifying cell phenotypes. Genome Biol 7, R100 (2006).

38. Berg, S. et al. ilastik: interactive machine learning for (bio)image analysis. Nat Methods 16, 1226–1232 (2019).

39. Schapiro, D. et al. histoCAT: analysis of cell phenotypes and interactions in multiplex image cytometry data. Nat Methods 14, 873–876 (2017).

